# Functional segregation of conversational production and comprehension when using word predictability

**DOI:** 10.1101/2024.06.18.599550

**Authors:** Caroline Arvidsson, Johanna Sundström, Julia Uddén

**Affiliations:** Stockholm University

**Keywords:** conversation, fMRI, surprisal, mPFC, language, theory of mind, cognitive control

## Abstract

The extent to which the language production and comprehension systems overlap remains debated. We address this debate using a dataset where participants engaged in unscripted conversations, while scanned with fMRI. Word predictability was hypothesized to rely on different processes, depending on whether the word was uttered or heard. We employed the information-theoretic measure of surprisal (the negative log probability of a word occurring, given the preceding context) as a parametric modulator, controlling for the word’s overall frequency. The results for production surprisal revealed activation in the left superior and inferior frontal gyri and motor areas. A large bilateral cluster in the posterior part of the medial prefrontal cortex extended from the supplementary motor area to the anterior cingulate cortex. The results for comprehension surprisal replicated findings from non-conversational contexts, showing involvement of the bilateral superior temporal gyrus/sulcus, presumably supporting bottom-up processes for prediction error detection. Importantly, no overlap in the neural infrastructure of production and comprehension was observed, suggesting that word predictability processes in production and comprehension differ. We suggest that while the comprehension system handles prediction errors, the production system minimizes these errors through adaptation, all to achieve successful communication.

## INTRODUCTION

The difference between language production and comprehension is currently disputed. Specifically, recent empirical papers debate the existence and extent of functional segregation of production and comprehension processes (Hu et al., 2022; Arvidsson et al., 2024; Giglio et al., 2022b; Matchin and Wood, 2020). In the case of functional segregation, one or more processes are more strongly, or even exclusively, recruited by one of the systems, given a particular situation. In comprehension, the prediction of upcoming linguistic input has been proposed to rely on production system processes (Dell and Chang, 2014; Pickering and Gambi, 2018). However, the accumulated evidence from neuroimaging does not favor the involvement of typical language production areas, such as the inferior frontal gyrus (LIFG), when listeners process unpredictable words (Willems et al. 2016; Lopopolo et al. 2017; Frank and Willems 2017; see however Russo et al. 2020). This raises the question of whether predictive mechanisms in comprehension are fundamentally dependent on production processes.

At the same time, knowledge about the underlying mechanisms of predictive processing in production is limited. Predictive processing in production may involve adaptive processes that are not required in comprehension. For example, speakers hyperarticulate and decrease articulation rate when producing words that are less predictable given context (Buz et al., 2014; Seyfarth et al., 2016), an effect that holds when processing ease is controlled for (Buz and Jaeger, 2016). In addition, word length (e.g., syllable length, number of syllables) positively correlates with a word’s average predictability in context (Piantadosi et al., 2009). These adaptations are potentially crucial to language processing in interactive settings, such as conversation.

In conversation, the listener predicts the incoming utterance (Levinson and Torreira, 2015), and can even begin planning their subsequent reply before their interlocutor has finished speaking (Magyari et al., 2014; Bögels et al., 2018; Bö gels, 2020). As predictability adaptations are only made in production, not comprehension, this indicates at least some segregation of production vs. comprehension. Whether this insight translates to segregation at the level of neural implementation has however never been tested. Presumably, any segregation is more detectable in conversation, where the pressure of meeting the communicative goals of language is increased. Therefore, the investigation of language processing during conversation is crucial in order to understand how production and comprehension processes may potentially diverge.

Previous fMRI studies have addressed the question of production-comprehension overlap, within and outside the perisylvian language network (Arvidsson et al., 2024; Giglio et al., 2022b; Hu et al., 2022; Matchin and Wood, 2020). This left-hemisphere dominant network of brain regions supports numerous linguistic (e.g., semantic, syntactic, and phonemic) processes. Most regions in this network are adjacent to the Sylvian fissure, including the LIFG, and the superior/middle temporal gyri (S/MTG).

Models on the neurobiology of language diverge regarding the extent of the production-comprehension overlap in the perisylvian language network. For example, Fedorenko et al. (2024) argue that language-specialized regions in this network support system-independent representations. These representations are computed in essentially two directions: encoding vs decoding. This contrasts with the updated version of the Dual stream model (Hickok, 2022). In this model, perisylvian network regions support some processes that are only required in production, such as the coding of linear sequences of morphemes, while all comprehension processes are shared with production.

In a previous fMRI study (Arvidsson et al., 2024), we investigated the neural overlap of the two systems. In contrast to Hickok (2022), and Fedorenko et al. (2024), we found that activation of the LIFG and the mPFC was stronger for production, while activation of the bilateral anterior S/MTG was stronger for comprehension. Stronger mPFC activation for production is particularly interesting for the current study, as this region has been associated with speaker adaptions. In Vanlangendonck et al. (2018), participants were required to adapt their utterances to the informational needs of the listener. These adaptations entailed choosing an appropriate adjective, based on the listener’s perspective. In the so-called communicative privileged ground condition, the listener’s perspective differed from that of the speaker. Production planning in the communicative privileged ground condition (vs. communicative linguistic and visual control) involved large portions of the bilateral dorsal mPFC, as well as the anterior mPFC. Whether these adaptive processes should be considered linguistic, or rather domain-general is unclear. Domain-general processes potentially involved in speaker adaptations are, for example, cognitive control/multiple demand processes, subserved by the multiple demand network Duncan (2010). A well-studied cognitive control component is working memory – the temporary storage and manipulation of the information necessary for complex cognitive tasks as language (Baddeley, 1992; Wager and Smith, 2003; Chai et al., 2018). It is also possible that speaker adaptations require processes involved in taking the listener’s perspective, i.e., theory of mind (Astington et al., 1988; Schurz et al., 2014).

Predictive processes operate on multiple levels of language (as discussed in, e.g., Kuperberg and Jaeger, 2016; Pickering and Gambi, 2018), including the word level. As an index of word predictability, it is common to use the information-theoretic measure of word surprisal. Surprisal is the negative log probability of a word occurring, given the preceding context (Hale, 2001).

Word predictability has been repeatedly investigated for comprehension of narratives in isolation, with the bilateral superior temporal cortex as a robust finding (Willems et al., 2016; Frank and Willems, 2017; Lopopolo et al., 2017; Russo et al., 2020). In the current study, we hypothesized that we would replicate these findings, when investigating conversational comprehension. The current study will be the first to involve a whole-brain investigation of the neural infrastructure of word predictability processes in production vs. comprehension (for an ROI analysis of differences in production vs. comprehension in a non-conversational context, see Giglio et al., 2022a). A pattern of partial segregation is expected when comparing word predictability processing in the two systems. Specifically, compared to comprehension, production may incur a greater cost on (1) processes in the LIFG, and (2) on adaptive processes, possibly located in the mPFC.

## METHODS

### Data

MRI images and orthographic transcriptions were obtained from a publicly available dataset by Rauchbauer et al. (2020). The current study included 24 participants (18 female, 6 male, M age = 28.8, SD = 12), excluding one from the original data set due to excessive head movement (max head movement > 4mm). In the Rauchbauer et al. (2020) data set, participants were scanned while they conversed in their L1 (French) with an interlocutor in the control room, via bidirectional audio and unidirectional video stream (the participant saw the interlocutor but not vice versa). The interlocutor was either an experimenter (who was said to also be a naïve participant) or a robot controlled by an experimenter. In the current study, we were not interested in production or comprehension events during the human-robot conversations. Therefore, we excluded these events by modeling them separately from the human-human events and only using the human-human events in the contrasts. Henceforth, all the mentioned events (i.e. production, comprehension, and silence events) refer to those during human-human conversations.

The MRI data in Rauchbauer et al. (2020) were collected with a 3T Siemens Prisma and a 20-channel head coil. Functional images used an EPI sequence with echo time (TE): 30 ms, repetition time (TR): 1205 ms, matrix size: 84 × 84, field of view (FOV): 210 mm × 210 mm, voxel size (VS): 2.5 × 2.5 × 2.5 mm^3^, 54 slices co-planar to the anterior/posterior commissure plane (axial), flip angle: 65 degrees, multiband acquisition (factor 3). Structural images were acquired with parameters TE: 0.002 ms, TR: 2.4 ms, FOV: 204.8 × 256 × 256 mm, VS: 0.8 × 0.8 × 0.8 mm^3^, 320 slices (sagittal).

### Conversational events

Rauchbauer et al. (2020) automatically segmented audio files into blocks of speech. The start and end of a block were defined as silence > 200 ms. The speech blocks were visually inspected and manually transcribed. In the present study, these blocks of speech were categorized as either a production or a comprehension event, depending on whether it was produced by the participant (production) or the experimenter (comprehension). Silences (i.e., periods during which both interlocutors were silent) were modeled as a separate event type. As the objective of the current study was to investigate word-level processing, we explored automated methods to segment the production and comprehension events into individual words. AI-based word segmenters (e.g., Whisper) did however not produce sufficiently accurate onset and offsets on our French data, the way we applied it. Instead, we approximated the onset and duration of individual words, based on the duration and number of words of each utterance. We calculated the (a) word duration and (b) word onset as follows:

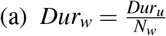

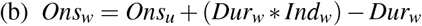

where *Dur*_*u*_ is the duration of the utterance and *Dur*_*w*_ is the word duration, *Ons*_*u*_ is the utterance onset and *Ons*_*w*_ is the word onset, *N*_*w*_ stands for the number of words in an utterance, and *Ind*_*w*_ stands for the word index (the first word has an index of +1 and the last word has the index of *N*_*w*_). Visual inspection ensured that this method was more reliable than using an automated word-segmenter, (e.g., Whisper). For an illustration of the conversational events, see **Figure 1. Table 1**. Shows descriptive statistics for the number of words per participant.

**Table 1.** Mean number of words in each event type (production, comprehension), per participant.

| Event type | Mean N words | Stdev N words |
| --- | --- | --- |
| Production | 910 | 299 |
| Comprehension | 996 | 258 |

**Figure 1.**
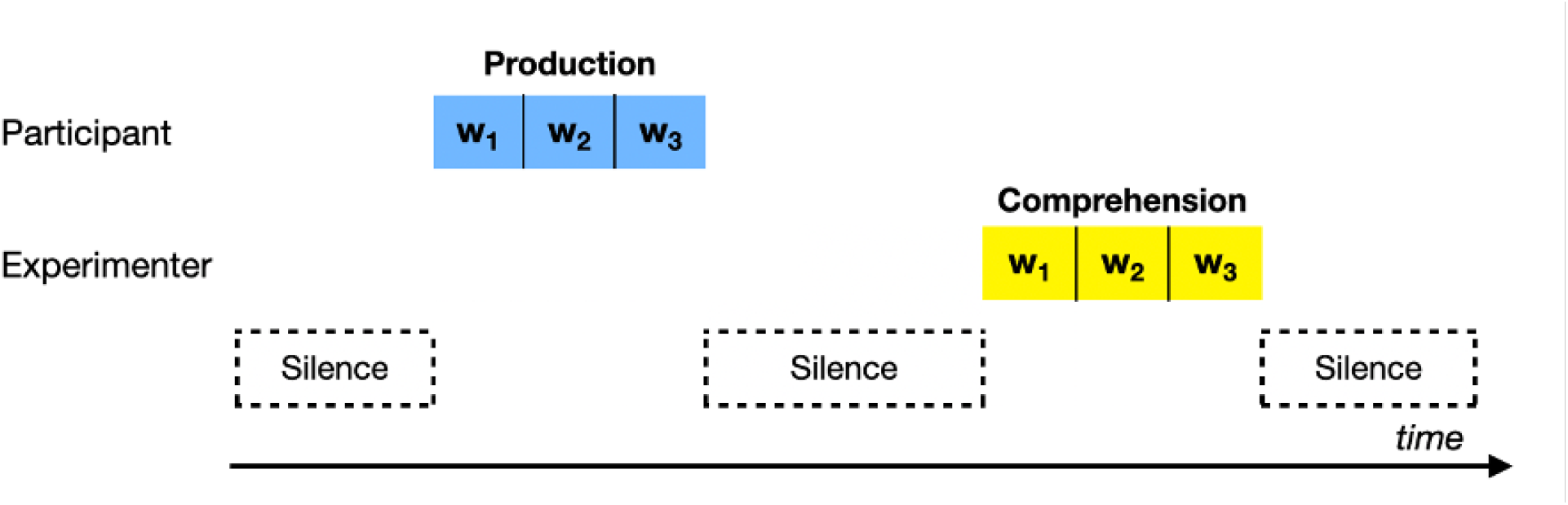
The conversational events production, comprehension, and silence. Production was speech produced by the participant. Comprehension was speech by the experimenter. Both production and comprehension events were segmented on the word level. Silences, e.g., between two speakers turns, were modeled as a separate event type.

### Ethical Statement

Ethical approval was obtained by Rauchbauer et al. (2020) from the ethics committee ‘Comité des Protection des Personnes Sud Mediterranneé I’ adhering to EU data protection laws.

### fMRI Preprocessing

Preprocessing was performed using fMRIPprep (v21.0.1, Esteban et al., 2019). The T1-weighted (T1w) image was skull-stripped and corrected for intensity non-uniformity, and furthermore used as reference throughout the workflow. Volume-based spatial normalization to standard space (MNI152NLin2009cAsym) was performed through nonlinear registration with ‘antsRegistration’ (Avants et al., 2009), using brain-extracted versions of both T1w reference and the T1w template.

The following preprocessing was performed for each of the BOLD runs per subject. Head-motion parameters (transformation matrices, and six corresponding rotation and translation parameters) were estimated. Spatiotemporal filtering was conducted using mcflirt (part of the FSL package Jenkinson et al., 2012). The BOLD time series was resampled onto native space by applying the transforms as an initial step to correct for head motion. The BOLD reference was then co-registered to the T1w reference. Co-registration was configured with six degrees of freedom. We included additional steps to correct for head motion. A set of physiological regressors was extracted to allow for component-based noise correction. The removal of non-steady state volumes and spatial smoothing with an isotropic, Gaussian kernel of 6mm FWHM (full-width half-maximum) was performed before automatically removing motion artifacts using independent component analysis (ICA-AROMA). Head movements in coordinates x, y, and z were inspected independently. As previously mentioned, one participant was excluded from further analyses because their max head movement exceeded 4 mm. The average framewise displacement (FD) across participants was 0.65 mm. None of the participants in the current data set exceeded an FD value of 2.5 mm, which has been used as an exclusion limit in previous studies (Giglio et al., 2022b). The preprocessed AROMA-cleaned images were used in the current analyses.

### Whole-brain analysis

For the first-level single-subject analysis, production, comprehension, and silences were modeled as three regressors. Images acquired during the presentation of the fixation cross (fixation) and the presentation of the advertising image (advertisement) were modeled as two separate regressors. Additionally, two covariates (called “parametric modulations” in SPM12) were added: word surprisal and ln-transformed (natural logarithm) word frequency. Word suprisal was the variable of interest, estimated using the GPT-2 language model fr-boris (Müller and Laurent, 2022) and the Python package Surprisal (Sathe, 2023). Unlike the trigram model used in Willems et al. (2016), the current study calculated word surprisal measures based on the entire utterance in which the word appeared (average length of 6.06 words). Moreover, log-transformed lexical frequency was obtained for each word using a French web corpus (Web 1T 5 gram, Brants and Franz, 2009) and was used as a control parameter to account for general word frequency. This meant that the column containing frequencies was placed before the surprisal column in the design matrix. A default value of 0 was assigned to words that were not found in the frequency list (constituting 2.6% of the data, or 1237 out of 47636 non-unique words). Head movements were modeled as six motion parameters. The events were convolved with a canonical hemodynamic response function. The effect of surprisal was computed in the first-level contrasts for production and comprehension respectively, by setting surprisal (in production or comprehension) to 1 and all other regressors to 0.

The second level analysis was conducted with one-sample t-tests on the contrast images defined at the first level. A cluster-forming threshold of an uncorrected *p*-value was set to .001 (no extent-level threshold, *k* = 0). Family-wise error (FWE), as implemented in SPM12, was used as the multiple comparison correction method (cluster and peak level). Only clusters with *p*_*FWE*_ < .05 at cluster level were reported in the current investigation. The test statistic of each cluster’s highest peak (voxel) is also reported. No additional voxels were reported, even if they were significant at *p*_*FWE*_ < 0.05 at the voxel level. Cluster labeling was performed automatically using the Automated anatomical labeling atlas 3 toolbox for SPM (AAL3, Rolls et al., 2020) and manually inspected in MRIcron using the Harvard-Oxford probabilistic atlas ‘HarvardOxford-cort-maxprob-thr0-1mm’.

### Testing for overlap in the mPFC

In a follow-up analysis, we wanted to investigate whether eventual mPFC activation in the two surprisal contrasts should be interpreted as relying on language, working memory (cognitive control), or theory of mind (ToM) processing. This was done by testing for the number of overlapping voxels with neurosynth association masks, using the search-terms “language”, “working memory”, and “tom”. We used an mPFC mask (as defined in the AAL3), to mask all images, before counting the number of overlapping voxels.

## RESULTS

### Whole-brain results

First, we tested whether there were voxels across the whole brain whose activation correlated positively with word surprisal (See **Figure 2** for a visualization; **Table 2** for cluster labels and inferential statistics). For conversational production, statistically significant activation was observed in six clusters. The largest cluster was found in midline structures of the frontal cortex, including the bilateral SMA and ACC. Three lateralized clusters were found in the motor cortex, two in the right and one in the left pre- and postcentral gyri, and adjacent structures, such as the LIFG (pars opercularis) and the left insula. One cluster was found in the occipital cortex bilaterally, inter alia including the superior occipital gyri and cuneus. The final cluster for the production surprisal vs. implicit baseline contrast was found in the cerebellum. For conversational comprehension, we observed two clusters of activation: one in the right and one in the left temporal lobe. The right temporal cluster was more strongly activated and extended more anteriorly than the left temporal cluster.

**Table 2.** Activations for the contrasts production suprisal >implicit baseline, and comprehension surprisal >implicit baseline. The number of voxels is given for each cluster, together with the MNI coordinates and t-value of the cluster’s maximum peak (local maxima).

| Anatomical region | Local maxima |  |  | Cluster |  | Voxel |  |
| --- | --- | --- | --- | --- | --- | --- | --- |
| | x | y | z | Size | $p_{FWE}$ | t-value | $p_{FWE}$ |
| *Production surprisal* |  |  |  |  |  |  |  |
| (L/R)SMA/(L/R)MCing/<br>LSFG/LSFG/(L/R)ACC | -6 | 4 | 58 | 1957 | <.001 | 9.41 | <.001 |
| RPreCG/RPostCG/RRolOper | 46 | -16 | 42 | 1237 | <.001 | 9.36 | <.001 |
| LPostCG/LPreCG/LInsula/<br>LIFG(oper)/LRolOper/LPutamen/ | -46 | -16 | 40 | 2151 | <.001 | 8.58 | .001 |
| RPreCG/RPoCG | 20 | -28 | 60 | 114 | .04 | 6.36 | n.s. |
| (L/R)Calcarine/(L/R)Cuneus/<br>(L/R)Lingual/(L/R)SOG | 16 | 84 | 6 | 1416 | <.001 | 5.99 | n.s. |
| Cerebellum | -14 | -62 | -20 | 143 | .02 | 5.26 | n.s. |
| *Comprehension surprisal* |  |  |  |  |  |  |  |
| RSTG/RMTG/RSTP/<br>RHeschl/RRolOper | 66 | -26 | 10 | 681 | <.001 | 6.53 | n.s. |
| LSTG/LMTG/LHeschl | -50 | -16 | 4 | 723 | <.001 | 6.41 | n.s. |

**Figure 2.**
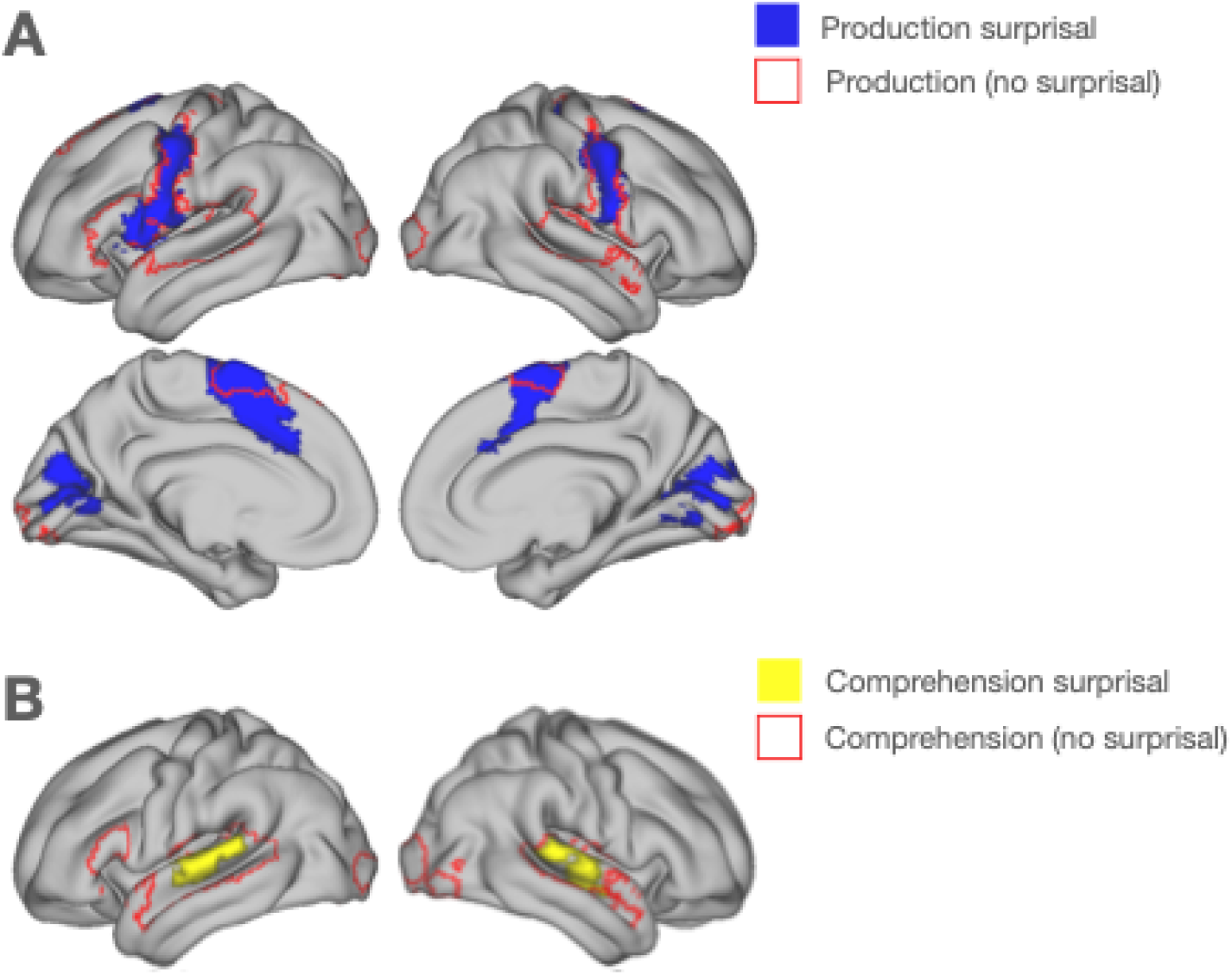
Whole-brain results showing regions reliably engaged with increased surprisal during conversational production (panel A, blue) and conversational comprehension (panel B, yellow). The red outlines show whole brain results observed in a previous study (Arvidsson et al., 2024, based on the same dataset) where we investigated the main effect of system (panel A: production, panel B: comprehension) without taking surprisal into account. The figure shows clusters with a cluster-forming threshold of puncorr = .001. Only clusters with a pFWE < .05 are reported.

### Overlap in the mPFC

We investigated the involvement of language, working memory (cognitive control), and theory of mind processes in the mPFC activation of our contrasts. This was done by counting how many voxels in a set of neurosynth activation maps were activated for our surprisal contrasts. For production, we found considerable overlap between production surprisal and two of the neurosynth masks: language (420 voxels in common) and cognitive control (464 voxels in common). There was no mentionable overlap between production activation and the theory of mind activation map (3 voxels). For an illustration of the overlap, see **Figure 3**. No mPFC activation was observed for comprehension surprisal.

**Figure 3.**
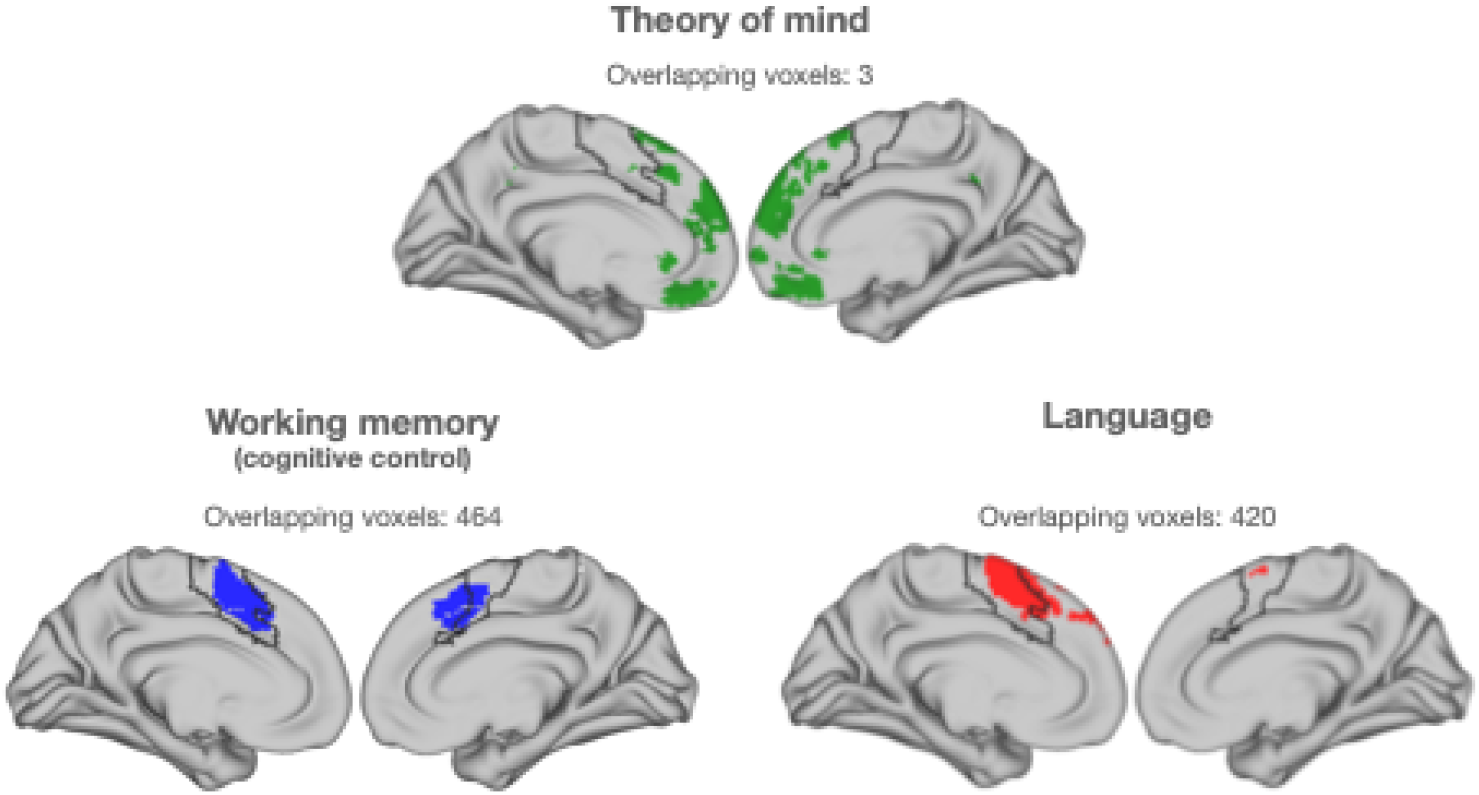
Overlap between production surprisal activation and the neurosynth activation maps. The black outline shows production surprisal activation. The activation maps were generated using the neurosynth default parameters, using the following keywords: “tom” (theory of mind); “working memory” (cognitive control); “language”.

## DISCUSSION

The current study has provided evidence of a segregation of conversational production and comprehension processes, in response to word predictability. Conversational production engaged lateral and midline structures of the frontal cortex, prominently within midline frontal structures such as the bilateral supplementary motor area (SMA) and anterior cingulate cortex (ACC). Additional activation was noted in lateralized clusters within the motor cortex, the left inferior frontal gyrus (LIFG, pars opercularis), and the occipital cortex. In contrast, conversational comprehension elicited activation in the bilateral superior temporal lobes. Further analysis of the medial prefrontal cortex (mPFC) revealed substantial overlap with neurosynth activation maps for language and cognitive control during production, suggesting a significant role of these cognitive processes in managing word predictability during speech production, whereas no such activation was found for comprehension. These findings underline the distinct neural mechanisms involved in processing language production and comprehension under conditions of varying word surprisal.

Word predictability processes in production entailed the involvement of regions in the mPFC. These results correspond to the activations generated by the communicative privileged ground condition in Vanlangendonck et al. (2018). These specific regions have previously been identified as supporting cognitive control and not theory of mind processing (see, e.g., the multiple demand network in Fedorenko et al., 2024; Duncan, 2010). As expected, the current evidence favors some involvement of domain-general cognitive control during word predictability processing in production. These findings need to be explained in light of previous empirical evidence, according to which predictive processing (although in comprehension) is domain-specific (i.e., selectively supported by the language network).

The results speak against the involvement of theory on mind in word predictability processing during production. This is hard to integrate with descriptions in which pragmatic processing is equated to theory of mind (Fedorenko et al., 2024; Wilson and Sperber, 2002), considering that speakers have been shown to make adaptations depending on how easily a word is anticipated, given context (Buz et al., 2014; Buz and Jaeger, 2016; Seyfarth et al., 2016).

In contrast to our earlier investigation of general production and comprehension processes (Arvidsson et al., 2024), the current study utilized a computational measure that allowed us to investigate specific processing aspects of the systems. Other studies have focused on how specific tasks differ in production and comprehension (e.g., phrase structure building, lexical access; Hu et al. 2022; Giglio et al. 2022b; Segaert et al. 2012; Segaert et al. 2013; Menenti et al. 2011). Yet, this approach has not been employed in whole brain investigations, using predictive processes.

In conclusion, our study provides evidence for a partial segregation of the neural mechanisms underlying word predictability in conversational production and comprehension. In conversational production, word predictability processing entails the recruitment of the left superior and inferior frontal gyri, motor areas, and the medial prefrontal cortex. In contrast, word predictability processing in conversational comprehension engages the the bilateral superior temporal gyrus/sulcus. The observed involvement of the superior temporal gyri is in line with previous research on word predictability in narrative comprehension (Willems et al., 2016; Frank and Willems, 2017; Lopopolo et al., 2017; Russo et al., 2020). The absence of overlapping infrastructure of word predictability processes in conversational production and comprehension highlight the specialized functions each system performs with respect to word predictability. In regard to word predictability, our findings indicate that the comprehension system focuses on detecting prediction errors, while the production system adapts to minimize these errors, facilitating effective communication.

